# Magic-BLAST, an accurate DNA and RNA-seq aligner for long and short reads

**DOI:** 10.1101/390013

**Authors:** Grzegorz M Boratyn, Jean Thierry-Mieg, Danielle Thierry-Mieg, Ben Busby, Thomas L Madden

## Abstract

Next-generation sequencing technologies can produce tens of millions of reads, often paired-end, from transcripts or genomes. But few programs can align RNA on the genome and accurately discover introns, especially with long reads. We introduce Magic-BLAST, a new aligner based on ideas from the Magic pipeline. It uses innovative techniques that include the optimization of a spliced alignment score and selective masking during seed selection. We evaluate the performance of Magic-BLAST to accurately map short or long sequences and its ability to discover introns on real RNA-seq data sets from PacBio, Roche and Illumina runs, and on six benchmarks, and compare it to other popular aligners. Additionally, we look at alignments of human idealized RefSeq mRNA sequences perfectly matching the genome. We show that Magic-BLAST is the best at intron discovery over a wide range of conditions and the best at mapping reads longer than 250 bases, from any platform. It is versatile and robust to high levels of mismatches or extreme base composition, and reasonably fast. It can align reads to a BLAST database or a FASTA file. It can accept a FASTQ file as input or automatically retrieve an accession from the SRA repository at the NCBI.

## INTRODUCTION

RNA-seq and DNA-seq experiments generate tens of millions of reads sampled from transcripts or genomes. The resulting data allows investigations that include, but are not limited to, gene expression, gene structure, nucleotide and structural variations. Different analysis approaches are available for some investigations. For example, coarse gene expression can be studied with alignments or with alignment-free methods (1). On the other hand, the investigation of fine grain gene structure does require alignments or sequence assembly and may benefit from a specific sequencing technology. While relatively short reads, 50 or 100 bases paired end Illumina style reads are sufficient for coarse gene expression profiling and most introns or single-nucleotide variant (SNV) calling, other studies such as SNV phasing, full length transcript description or structural genomic rearrangements are facilitated by longer reads. But not all aligners can handle longer reads. Towards comprehensive and reliable mapping and variant discovery, the aligner should provide good tolerance to mismatches and robust discrimination of reads mapping ambiguously at multiple quasi-repeated sites. Several groups have written fast aligners (2,3), but a recent benchmark by Baruzzo and the Grant team showed that new aligners could still offer value to the community (4).

Here, we present Magic-BLAST, the RNA-seq splice aware and pair aware new member of the BLAST family of programs. We compare Magic-BLAST to three other popular aligners. We look at a variety of RNA-seq read lengths, between 100 bases and 100 kilobases, from Illumina, Roche 454 and PacBio as well as full length RefSeq transcripts and find that most aligners can accurately map short Illumina-style human paired reads with few mismatches, but that aligners do not work as well with longer reads. We also find that higher numbers of mismatches as well as compositionally biased genomes pose problems for some aligners. Magic-BLAST can handle the different sequencing technologies, error rates, and compositional bias without special tuning, and this should allow, in particular, to map an RNA-seq experiment to the genome of a related species when a good quality reference genome is not available. Additionally, Magic-BLAST outperforms all other aligners in mapping of long reads and in identification of introns, including discovery of unannotated introns, under all circumstances, even though it is single pass and is aligning RNA to the genome without knowledge of a transcriptome.

We chose the name Magic-BLAST to emphasize the ideas and software that went into the tool. Magic-BLAST derives its core algorithms from the Magic pipeline, described in detail in the supplementary material of (5). It is implemented using the same C++ framework as the BLAST+ executables (6,7,8). The merger of these tools results in a versatile and robust aligner. Magic-BLAST implements the Magic ideas of checking for overrepresented target sequence fragments during seed selection as well as an innovative greedy alignment extension procedure. Magic-BLAST can produce spliced alignments when aligning RNA on genome and selects the best-matching position genome wide by optimizing a spliced alignment score for single and for paired reads. Magic-BLAST has a more limited role than its namesake. Magic refers to an entire pipeline with high level functionalities, but Magic-BLAST implements just the Magic aligner.

Magic-BLAST takes advantage of the existing BLAST infrastructure. It aligns reads against a BLAST database, which is the same database used by other BLAST programs. It can also align reads against a FASTA file or even just an accession, with the actual sequences automatically retrieved from the NCBI. Sequence reads can be provided as SRA accessions or as files in FASTA or FASTQ format. Magic-BLAST can transparently gzip or gunzip the sequence reads and/or the reference FASTA or FASTQ files. It was field-tested in several NCBI hackathons that provided substantial feedback on features and usability.

We compare Magic-BLAST to three popular aligners, HISAT2 (9), STAR (10,11), and TopHat2 (12, 13), also evaluated in (1) and (4). We look at precision (percentage of results that are correct), recall (percentage of the total true positives returned) and F-score (harmonic mean of recall and precision) for alignments and for intron discovery, as measured on truth-bearing RefSeq (14) human transcripts (assembled to exactly match the genome), on experimental long or short reads, and on simulated benchmark data assessing the impact on the alignments of variable levels of polymorphisms and errors, up to mimicking an alignment to a related species (4). The last data set additionally tests the aligners on a genome with extremely biased base composition, using the malaria agent *Plasmodium falciparum*, which is 80.7% AT. Magic-BLAST is not the fastest tool but is reasonably fast and works well when mapping RNA to the genome or to the transcriptome. For RNA-seq, it auto-adapts to the type of data and read length without requiring from the user any choice of parameters. It is versatile, easy to use, robustly precise and conservative in all circumstances.

## MATERIAL AND METHODS

### Algorithm Overview

The Magic-BLAST algorithm has a structure similar to that of other BLAST programs (8). It reads the data in batches and builds a “lookup table”, which is an index of word locations in the reads, 16-bases by default. It then scans the database sequences, usually a reference genome, for matches in the lookup table and attempts to extend selected initial matches to the length specified by the user (18 by default). The resulting matches form a seed for computation of local gapped alignments. Collinear local alignments are combined into spliced alignments. Exons shorter than the seed length cannot be captured, but they are rare (less than 0.2% of RefSeq introns), and most will be recognized by aligning in parallel on the known transcriptome. For paired reads, the best alignments are selected based on the alignment quality of the pair. For example, if one read of a pair maps equally well at two genomic sites, and the second read maps best at a single site, the read pair will be reported as mapping uniquely at the position dictated by the second read. In this way, the specificity of the mapping truly reflects the number of bases sequenced in the whole fragment, i.e. 200 bases specificity for 100+100 paired-end reads. Below, we present a detailed description of the above steps.

### Repeat filtering

Most genomes contain interspersed repeats and gene families that complicate correct placement of reads in a genome. To avoid seeding to ambiguous positions, Magic-BLAST scans the reference sequence and counts 16-base words. Those words that appear in the reference database more than a user-specified number of times (by default 60) are not indexed in the lookup table, so that they never form a seed alignment. To make this procedure more efficient, only words present in the reads are counted. The cut-off number 60 was selected experimentally as the best trade-off between sensitivity and runtime for RNA-seq. Additionally, Magic-BLAST specifically masks out 16-base words that contain at least 15 A’s or 15 T’s, effectively avoiding seeding on poly-A tails. This approach is similar to soft masking in other BLAST programs.

### Local gapped alignment

Magic-BLAST computes a local alignment by extending exact word matches (18-bases by default) between a read and a reference sequence. We use a simplified greedy alignment extension procedure, previously used in Magic (5). Starting with the seed, the alignment is extended until the first mismatch. Next, we attempt to characterize the mismatch as a substitution, insertion or deletion of one to three bases by recursively testing the quality of the alignment of the following few bases. This is done by applying successively a table of candidate alignment operations (Table 1) until the associated requirement is met. A requirement is that a specific number of bases must match within a given number of bases following the applied operation. The first operation whose requirement is met is applied to the alignment and the algorithm proceeds to the next position on both sequences. A single substitution is reported if no requirement is satisfied. The list of alignment operations and their associated conditions used in Magic-BLAST is presented in Table 1.

**Table 1.** List of alignment operations used in the Magic-BLAST alignment extension.

| Operation | Length | Required number of matching bases | Allowed number of mismatches |
| --- | --- | --- | --- |
| Substitution | 1 | 9 | 0 |
| Insertion | 1 | 10 | 0 |
| Deletion | 1 | 10 | 0 |
| Insertion | 2 | 10 | 0 |
| Deletion | 2 | 10 | 0 |
| Insertion | 3 | 13 | 0 |
| Deletion | 3 | 13 | 0 |
| Substitution | 2 | 12 | 0 |
| Insertion | 1 | 10 | 2 |
| Deletion | 1 | 10 | 2 |
| Insertion | 2 | 10 | 2 |
| Deletion | 2 | 10 | 2 |
| Insertion | 3 | 13 | 2 |
| Deletion | 3 | 13 | 2 |

Figure1 shows an example alignment extension. First, there are two matches and the algorithm moves to the right by two positions on both sequences. When a mismatch (T-G) is encountered the algorithm tries successively each alignment operation and checks its requirements. The first operation, a substitution which requires nine matching bases following the mismatch, fails. The second operation, an insertion which requires ten consecutive matches, succeeds and is applied to the alignment. In the last step there is a match (G-G).

**Figure 1.**
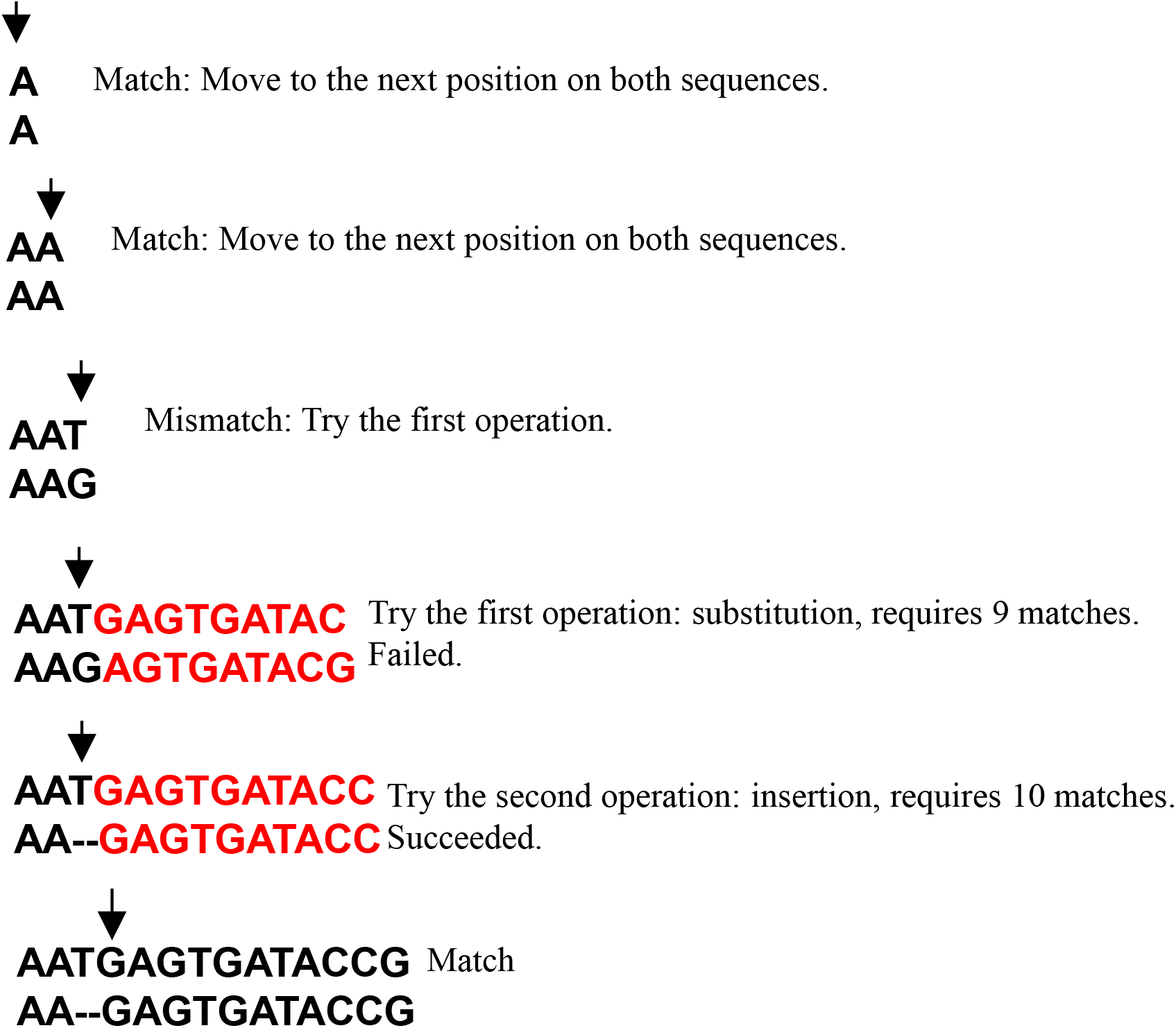
An example alignment extension procedure. The arrows point at each step to the position currently considered in both sequences. First, there are two matches and the arrows move to the right by two positions on both sequences. When a mismatch (T-G) is encountered the algorithm tries alignment operations and conditions. The first operation: a substitution, which requires nine matching bases following the mismatch, fails. The second operation, an insertion which requires ten matches succeeds and is applied to the alignment. In the last step there is a match (G-G).

We use the X-drop algorithm (6) to stop the extension. At each position, we record the running alignment score. The algorithm stops at the end of a sequence or when the current score is smaller than the best score found so far by more than the most penalized gapped alignment operation (three-base gap in Table 1). The algorithm then backtracks to the position with the best score.

Because most reads align to a reference with few or no mismatches, this method is faster and more memory efficient than the dynamic programming-based extension procedure used in other BLAST programs. Moreover, this approach facilitates collection of traceback information at little additional cost. This method can be tuned to a given sequencing technology for an expected rate of mismatches or gaps simply by adapting Table 1. For example, in Roche 454 or PacBio, where insertions and deletions are more frequent than substitutions, one could switch to a modified table.

We compute an alignment score using the following system: 1 for each matching pair of bases, -4 for a base substitution, zero for gap opening (either a read or reference sequence), and -4 for each base of gap extension (insertion or deletion). A user can modify the mismatch and gap penalties. The quality coefficients present in the FASTQ file have no impact on the alignment score, and are not exported in the SAM output.

About half the time, a matching base can be placed on either side of a gap, so the gap can slide at equal score. To avoid difficulties in SNP detection, Magic-BLAST by convention shifts the sliding bases upstream of the gap, in the orientation of the target.

### Spliced alignments

Spliced alignments are found by combining collinear local alignments on a read and a reference sequence. Magic-BLAST constructs a chain of local alignments that maximizes the combined alignment score. It then updates the alignment extents so that the spliced alignment is continuous on the read and the intron donor and acceptor sites are, whenever possible, consistent with the canonical splice signals.

If two local alignments are continuous on a read (possibly with an overlap), then we first search for the canonical splice site (GT-AG or CT-AC) where the alignments meet. If this site is not found and each alignment has a score of at least 50, we search successively for the minor splice sites or their complement: GC-AG or CT-GC, AT-AC or GT-AT, then for any other non-canonical site. The first site found is accepted. The alignment score threshold of 50 was chosen because minor and non-canonical splice sites are rare, but pairs of di-nucleotides are frequently found in the genome. As a result, for reads shorter than 100 bases, Magic-BLAST conservatively only calls GT-AG introns.

Magic-BLAST also attempts to produce spliced alignments if a read has several local alignments separated by one to ten unaligned bases. First, we look for a splice signal within four bases of the end of the left alignment and, if found, we fix the beginning of the candidate intron. Second, we search for the corresponding end of intron signal at offsets that ensure a continuous alignment on the read, allowing at most one additional insertion or deletion. If this fails, the procedure is repeated with the end of the intron fixed and a search for the signal indicating the start of the intron. When the candidate splice signals are found, the alignments are trimmed or extended to the splice signals. The benefit of this method is that it correctly identifies introns even in the presence of a substitution, insertion, or deletion close to the intron boundaries. Because this procedure is very sensitive and can produce many spurious alignments, Magic-BLAST only allows the GT-AG signal in this situation.

The spliced alignment is scored with the same scoring system as the local alignment. There is no reward or penalty for splice sites and no preference is given to continuous versus spliced alignments. When mapping RNA to the genome, Magic-BLAST does not use an annotation file or a two-pass method. We recommend instead to map in parallel on the genome and on an annotated transcriptome, then use the universal scoring system of Magic-BLAST to select the best alignment, be it genomic or transcriptomic, for each fragment.

### Output

Magic-BLAST returns results in the Sequence Alignment/Map SAM/BAM format (15) or in a tab-delimited format similar to the tabular format in other BLAST programs, which is less standard but richer and easier to mine.

## RESULTS

### Datasets and programs

The ability of Magic-BLAST and other popular programs to map RNA-seq to genomes in a naïve fashion, without knowledge of an annotated transcriptome, and to find introns and their precise splice sites was assessed using seven truth-bearing datasets, one of which is new, and several experimental runs, from Illumina, Roche 454 and PacBio.

The new benchmark, called iRefSeq, is the image of the Human RefSeq mRNAs (14) exactly matching the genome. We selected the protein-coding RefSeq mRNAs, limiting to the 45,108 NM accessions that map to the main chromosomes and mitochondrial DNA of GRCh38 (GCF_000001405.36). These mRNAs are transcribed from 19,269 protein coding genes. Using the mapping coordinates, as given in the RefSeq annotation, we assembled genomic sequences into transcript sequences so that they exactly match the genome (see supplementary section 2.1). iRefSeq mRNAs range in length from 147 bases to 109,224 bases. This perfect data set forms a useful benchmark for RNA-seq aligners, because it seems simple to align, and each mismatch in the alignment indicates an imperfect mapping. Furthermore, the coordinates of the 210,509 distinct introns in iRefSeq are known.

The Baruzzo benchmark set of simulated RNA-seq reads, presented in (4), was also used. This set has some qualities that make it appealing for our analysis. The authors document their procedure well, they produce 10 million paired-end 100+100 Illumina-like reads at three vastly different error rates, nominally from 6.1 to 55 mismatches per kb, and they produce data for human and *Plasmodium falciparum*, a protozoan causing malaria in human (we refer to the latter sets as ‘malaria’). Baruzzo names these three different error rates T1, T2, and T3. The variable error rates simulate how the aligners would perform if the genome of the same species was available (T1), if only a poor-quality version of the genome or the sequence data was available (T2), and if only the genome of a related organism was available (T3). The malaria sets allow an analysis of how the aligners perform under extreme genome base composition as the genome is 80.7% AT. The human and malaria sets have the same number of reads, so the malaria sets are at least 100 times deeper, a confounding effect to unravel. In practice, each set is provided as triplicate benchmark runs, but since the results are very similar, only the results of R1 are shown in the figures (all 18 datasets are shown in Supplementary Tables S3, S4, S6). Surprisingly, we noticed that the measured level of mismatches per kb differs between the human and malaria sets: T1 has 5.4 and 6.5 mismatches per kilobase aligned in human and malaria respectively, T2 11.86 and 16.6, and T3 60.2 and 86.5.

Furthermore, we selected experimental RNA-seq data sets from the public Sequence Read Archive (SRA at NCBI) to represent three sequencing platforms, PacBio (SRR5009494, with 8285 reads of average length 1854 bases, sequenced from colon carcinoma cells), Roche 454 Titanium (SRR899420, with 416,045 reads of average length 348 bases, sequenced from the MAQC/SEQC brain mRNA sample (16)), and a deep Illumina HiSeq run (SRR534301, with close to 109 million 101+101 bases paired-end reads sequenced from fetal lung, also tested in (9)). Here, we will refer to each of these RNA-seq sets by its technology: PacBio, Roche and Illumina. Figure 2 presents a histogram of read lengths for the iRefSeq and these three experimental sets. To refine the conclusions, we later added two large PacBio runs (testes SRR5189667 and brain SRR5189652 from the GENCODE project (17), with non-native sequences edited using Illumina runs), two Illumina runs with longer read pairs (250+250 bases SRR5438850 from metastatic melanoma; 300+300 bases SRR5437876 from MCF7 cells, also studied in (1)). We finally truncated the Baruzzo human T1 sample to create a shorter 50+50 bases run and subsampled to 1% the malaria T1 sample to obtain, by keeping only every hundredth sequence, a coverage depth similar to that in human.

**Figure 2:**
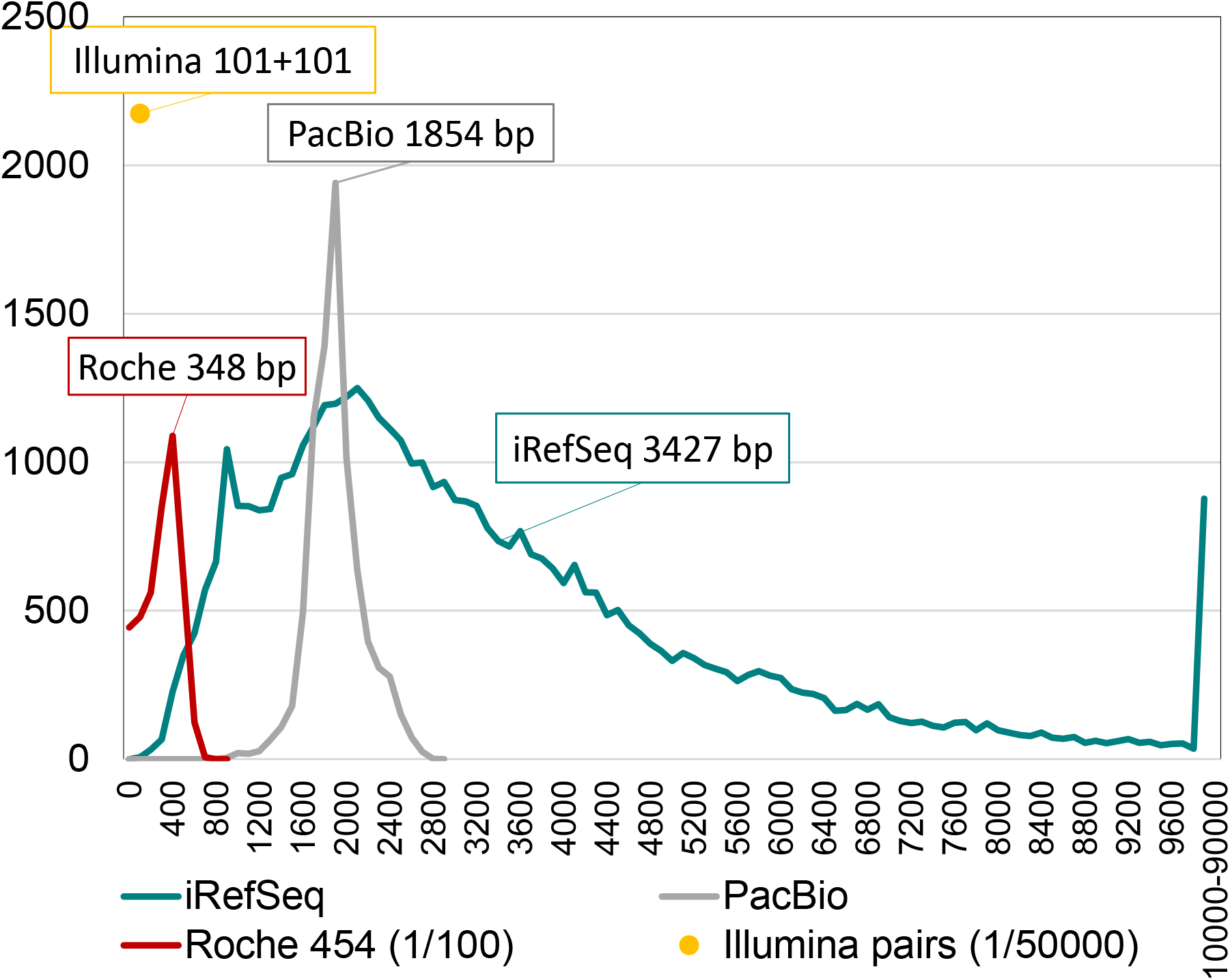
Actual read length histograms for the three experimental sets as well as iRefSeq. All Illumina reads have the same length (101+101 bases, paired-end) but other sets have non-trivial length distributions. Newer sequencing technologies tend to produce longer sequences. PacBio has the longest reads of the experimental sets, ranging from 710 to 2703 bases (average 1854 bases). Roche 454 Titanium reads span from 33 to 808 bases, with an average of 348 bases. iRefSeq has the longest reads of all sets: these full-length mRNAs range in length from 147 bases to 109,224 bases, with an average of 3427 bases. 9900 are longer than 10 kb. The Y-axis presents the number of reads per 100-base bin (on the X-axis). The Y scale is reduced 100 times for Roche 454, and 50,000 times for Illumina, which has the highest throughput of all technologies.

We examined the performance of several programs aligning RNA to the genome in the absence of a transcriptome (Human genome GRCh38 and *P. falciparum* genome provided in (4)). Magic-BLAST was compared to programs from 2009 to 2015: HISAT2 (9), STAR (10,11), STAR long (10,1,17) and TopHat2 (12, 13) (Details in supplementary section 1.2). The standard STAR is optimized for Illumina-type reads while the authors recommend STAR long for reads longer than 300 bases, but we tried both versions on all runs. The two-pass mode is recommended if no transcriptome annotation is provided: STAR long was run only in two-pass mode, but both one-pass and two-pass modes were tried for standard STAR. HISAT2 was run with default parameters as well as in a ‘relaxed’ mode which is more sensitive but much slower: the HISAT2 default parameters left 4,663 iRefSeq unmapped while all aligned in relaxed mode. Magic-BLAST and TopHat2 do not have a two-pass mode and were run with default parameters.

Two analysis programs were used in this project (Supplementary section 1): AliQC.py was developed in collaboration with Joe Meehan from the FDA for the SEQC project (16). It extracts, by comparing the SAM file to the genome, a detailed quality control on alignments, their length and multiplicity, and mismatches by reads, by type and by position along the read (i.e. by sequencing cycle) (Supplementary section 7). In case of multiple alignments, only the ‘primary’ alignment is considered in this analysis. The number of mismatches were confirmed using the NM: optional tag present in the BAM files. Another program, sam2gold.c, was written in C to compare the SAM files to the format in which Baruzzo (4) provided the benchmark truth (Supplementary sections 2 to 6). A master-script, described in Supplementary section 1.5 and deposited in GitHub (https://github.com/ncbi/magicblast), can download all the data, realign the sequences, and reproduce supplementary tables and sections 2 to 7, which support our entire analysis.

We first measure how well the different aligners identify introns, then we examine the properties of the alignments.

### Intron discovery

To test how well the aligners discover introns, the splice sites were extracted from the BAM output using the ‘N’ operation in the CIGAR string (15). For the iRefSeq and Baruzzo benchmark sets, the true position of each intron is known. The experimental sets do not come with such a “Ground Truth”, but a proxy for true and false positives are the introns annotated and not-annotated in iRefSeq. Of course, this is not strictly correct as some unannotated introns are no doubt real and just have not been discovered or annotated on the genome. On the other hand, it seems likely that all (or almost all) the annotated introns are real. This strategy allows a comparison of the results of all programs on all datasets and the measurement of precision, recall and F-score for intron discovery.

Magic-BLAST, HISAT2 in relaxed or standard mode, and STAR long are able to align very long reads on the genome and find introns. TopHat2 and the standard STAR failed to produce any results for very long reads, although TopHat marginally worked for Roche 454.

We first use a ROC curve approach (18) to precisely judge the quality of intron discovery, true versus false, as a function of minimal read coverage (Figure 3 and S3). We group the introns by read coverage in up to 100 bins. For each bin, the number of true positive (or annotated) introns are plotted on the Y-axis while the number of false positives (or unannotated) introns are on the X-axis. The resulting curves have up to 100 points and give us visual insight into how the different programs behave when the support, given by the number of reads mapped to each junction, increases. The truth for the benchmark sets is vertical and dark blue. The best curve, of course, would have all the true positives before any false positives, meaning the steeper the slope, and ultimately the higher to the left the curve is, the better. In fifteen cases, Magic-BLAST (red) is to the left and above all other curves and qualifies as the best intron finder in all conditions tested, for reads from 100 bases to 100 kilobases, with any level of mismatches, from perfect match to 8.6% mismatch. This observation applies to benchmark as well as all seven real data sets tested from PacBio, Roche or Illumina.

**Figure 3:**
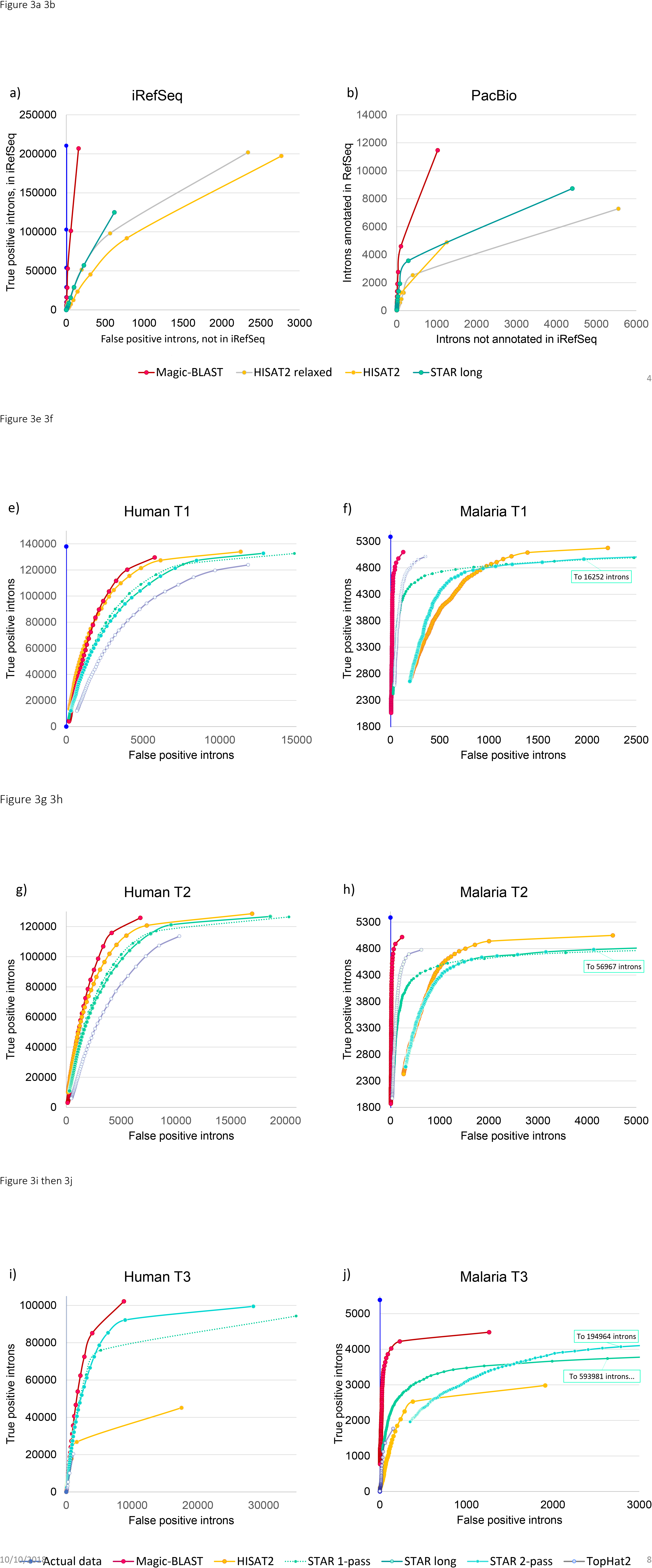
ROC curves showing intron discovery as a function of minimal read coverage, from 1 to 100 (or to the maximal observed coverage). The point with coverage 1, corresponding to all introns found in the experiment, is the point closest to the top right corner. The plots show, for each minimal coverage, the true positives on the Y-axis and the false positive on the X-axis. In the experimental sets, annotated and unannotated introns are used as a proxy for true and false positives. The best curve would have all the true positives before any false positives, meaning the steeper the slope the better. The benchmark sets, iRefSeq and six Baruzzo, have a built-in truth (vertical blue line in some graphs). a) For the iRefSeq set, because of the alternative splice variants, the truth has introns supported by 1, 2, and up to 51 RefSeqs, and Magic-BLAST (red) follows the truth remarkably closely, point by point. It finds slightly more true-positive introns than the HISAT2 programs, but the biggest difference is that HISAT2 finds fifteen to seventeen times as many false positive introns. STAR long finds only 60% of the introns with some false positives. b) For PacBio, with less than 9000 reads, Magic-BLAST already finds 11464 annotated introns, many more than the two HISAT2 versions and STAR long, yet it finds the fewest unannotated introns. c) The Roche presents a similar, though less extreme, result. d) In Illumina (zoomed in supplementary figure 3.1), Magic-BLAST followed by STAR 1-pass have the steepest slopes. Then come STAR 2-pass and TopHat2, then HISAT2; these last three aligners call unannotated introns at high coverage. e) to j) In the Baruzzo shallow human T1 (e) and T2 (g) benchmarks, Magic-BLAST then HISAT2 perform the best, followed by STAR 1- then 2-pass, then TopHat2. i) In human T3, HISAT2 and TopHat drop considerably, only Magic-BLAST and STAR can find introns in the presence of a high level of mismatches. f) h) j) In the ultra-deep malaria sets, Magic-BLAST remains best, STAR 2-pass and HISAT2 drop below TopHat. At coverage 1, STAR has by far the largest number of false positives (Supplementary Figures S3).

This observation can be summarized by quantifying intron precision, recall, and F-score for the iRefSeq and experimental sets (Figure 4a-c and S4). For the experimental sets where we use RefSeq annotated introns as truth, only the comparison of the scores of the various aligners is meaningful: on the same data, they should detect the same introns. But the precision and the recall depend on the tissue and on the depth of the experiment. The Illumina 101+101 run is from fetal lung, a stage not well represented in the RefSeq collection, and this explains the apparently low precision (below 46%): many of the observed introns are probably real but not annotated. At the same time, RefSeq annotates introns and genes from all tissues, and typically a sample derived from a single tissue and sequenced deeply will express 70 to 75% of all annotated genes and introns, this explains the recall around 72% in this Illumina set. For the shallow data sets from Roche adult Brain and PacBio colon carcinoma cells, precision seems good (up to 92% in Magic-BLAST) because these tissues were used intensely in RefSeq annotation, and because at low coverage, one sees mainly the highly expressed genes, which are the best annotated. Yet the low depth of sequencing explains why Roche finds only 31% of the annotated iRefSeq introns, and the very small PacBio run finds less than 6%.

**Figure 4.**
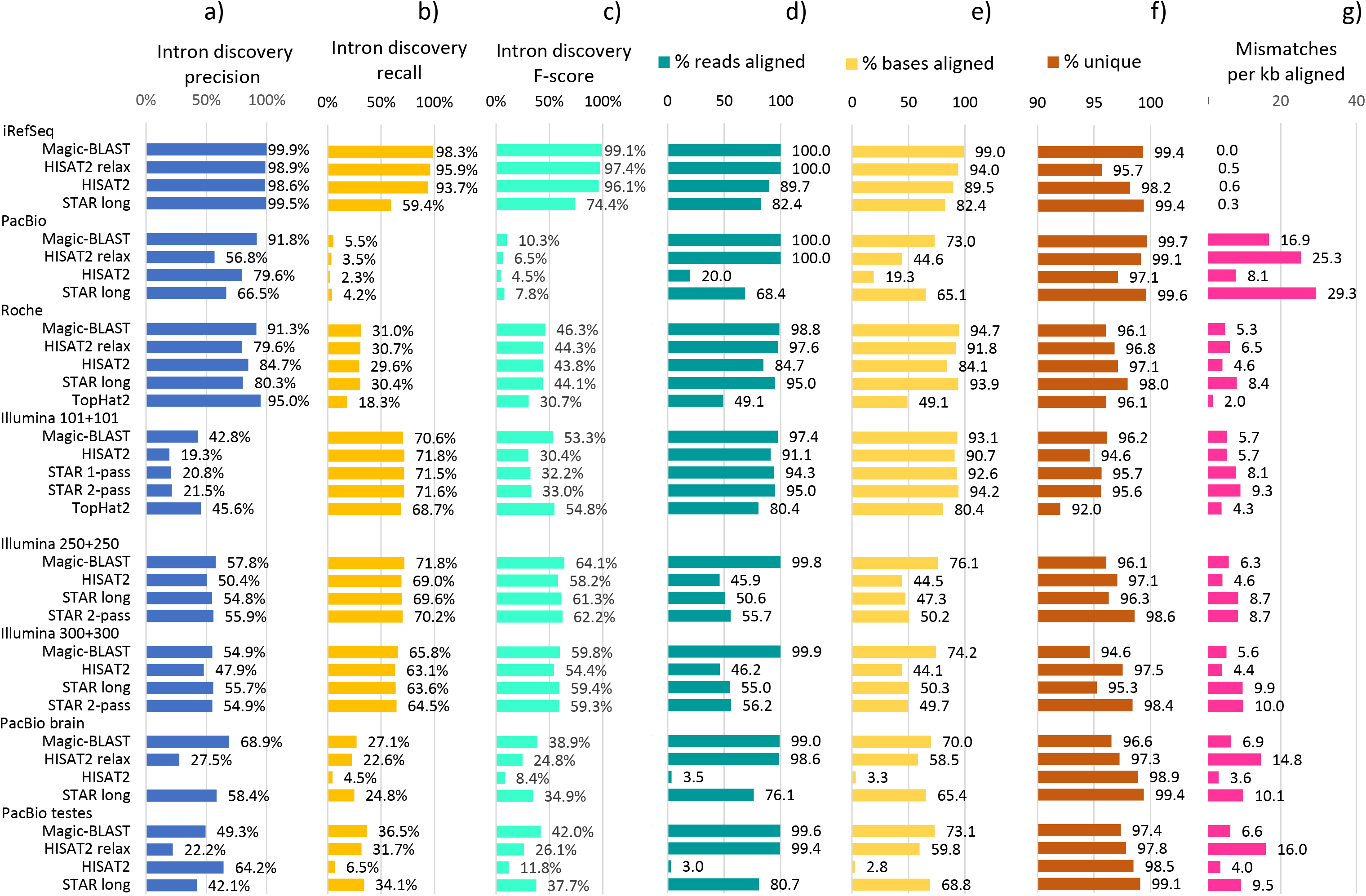
Performance of the aligners on intron discovery and alignment statistics measured on iRefSeq and seven experimental datasets. a) Intron discovery precision p=(TP/TP+FP). b) recall r=(TP/TP+FN) and c) F-score (2pr/p+r). All introns are counted, including those with a single support. An intron annotated in iRefSeq is counted as true, an unannotated intron as false. d) percentage of reads aligned. e) percentage of bases aligned. f) percentage of reads aligned that map to a single genomic site (uniquely mapped). The read length varies along the plot. Average sequenced length per fragment are: iRefSeq 3427 bases, PacBio 1854 bases, Roche 348 bases, Illumina 202, 500 and 600 bases, PacBio 1300 and 1323 bases. The longer the reads, the more specific their alignments are. g) number of mismatches per kb aligned. This number reflects at the same time the accuracy of the sequencer and the capacity of each aligner to cope with the mismatches and to perform accurate mapping.

**Figure 5.**
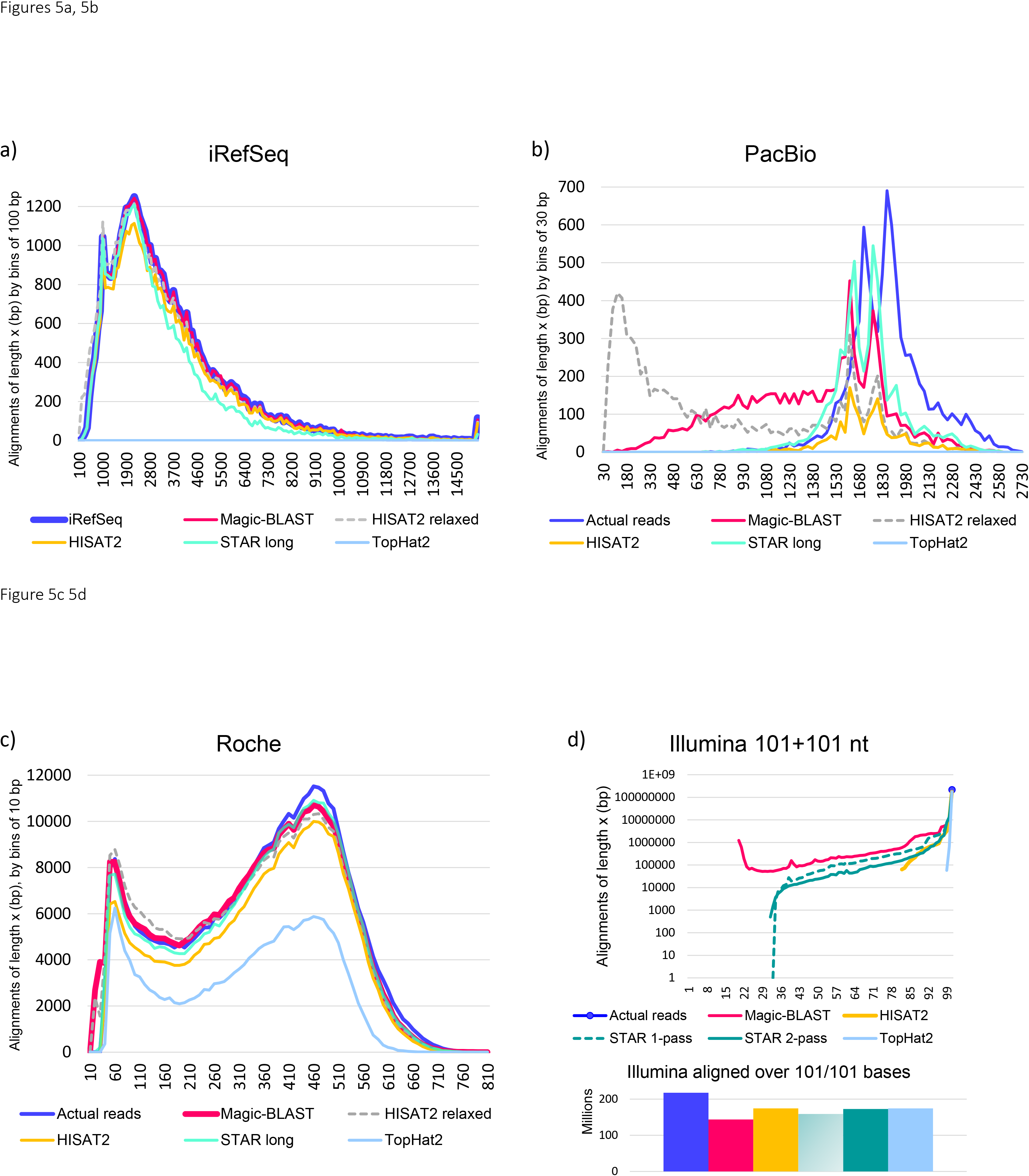
Histograms of read and alignment lengths for iRefSeq (a), PacBio (b), Roche 454 (c) and Illumina (d) data sets. The X-axis shows the length in bases, the Y-axis the number of reads in each length bin, in the original dataset (blue) or after alignment. The counts are binned every 100 bases for iRefSeq, every 30 bases for PacBio, every 10 bases for Roche and every base for Illumina. The Y scale for Illumina is logarithmic; a complementary zoom on the point at 101 bases, which represents the fully aligned reads, is shown below as a bar plot.

The ROC curves make it apparent that the ability of the aligners to discover introns, with a good balance of true to false positives, changes as the read coverage for introns decreases. We use this insight to calculate a coverage dependent best F-score for the aligners (Supplementary sections 3 and 4). At the best F-score, Magic-BLAST has the highest F-score in 15 experiments, plus the 12 duplicate benchmarks. The only exception is the human T1 truncated at 50+50, where HISAT2 takes the lead. Magic-BLAST also reaches its best score at the lowest coverage of all aligners in almost all cases. The other aligners achieve optimal scores at much higher coverage than Magic-BLAST for the deep experimental Illumina and Baruzzo malaria sets. Magic-BLAST is more conservative than the other programs, and even the introns supported by a single read appear reasonably trustworthy.

Another notable feature in the ROC curves for the Baruzzo benchmark is that STAR produces the largest number of false positives in every case, followed by HISAT2 (Figure 3, Supplementary section 3). In the deep malaria set, which has about 5500 annotated introns, STAR produces up to 600,000 false positive introns (Figure 3j). This greedy intron-finding behavior fits with the observation that introns and splice sites found by STAR and HISAT2 at low coverage are mainly untrustworthy. It is worth noting that, as judged by the ROC curves, HISAT2 produces much worse results in the very deep malaria T1 than in the shallow human T1 (Figure 3e and 3f), and that STAR 2-pass can produce worse results than the 1-pass version. This is especially apparent for deep sets such as the experimental Illumina and the Baruzzo malaria (Figures 3fhj, Supplementary figure S3.1). This hypothesis was tested and confirmed by subsampling 1% of malaria T1: STAR 2-pass became just as good as STAR 1-pass (compare Figure 3f and Figure S3.3) and the false positive introns went away much faster than the true positives in STAR and HISAT2.

Figure 7g presents intron discovery F-scores for the Baruzzo set. Magic-BLAST has the best F-scores for all the sets, but really excels for the T3 human set and all the malaria sets.

### Alignment quality

Various metrics can be applied to characterize the ability of aligners to accurately map the reads. For simulated runs, where a truth is known by construction, one can compare the precise placement of each alignment to the ‘true’ position of the read and partition all alignments in one of four categories: completely correctly aligned (True positive type 1), partially but correctly aligned (True positive type 2), misaligned (False positive) or not aligned (False negative). This provides a direct measure of alignment precision, recall and F-score (Figures 6c, 7a-c, S6.1).

**Figure 6.**
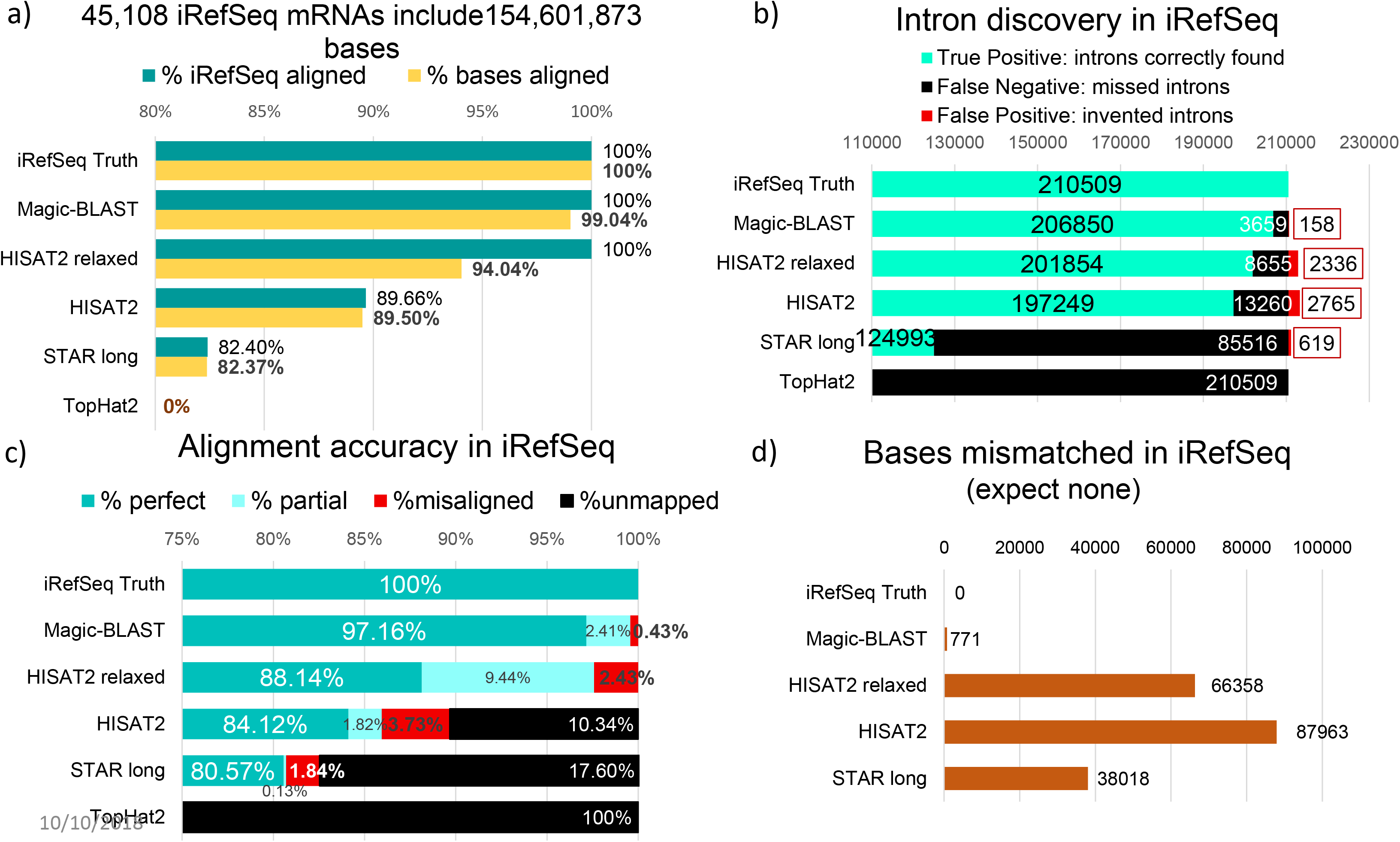
Characteristics of alignments of the iRefSeq set. a) The 45,108 iRefSeq include 154,601,873 bases exactly matching the genome (Truth). For each program, the percentage of iRefSeq sequences aligned (green) and the percentage of bases aligned (yellow) are given. In case of multiple alignments, each read contributes only once, at its primary position. b.) Intron discovery in iRefSeq: the number of introns correctly found (green, TP), the number present in the truth but missed by the aligner (black, FN) and the number of invented introns, found by the aligner but absent from the truth (red, FP) are shown. c) Accuracy of the alignments: the true alignment of each read is defined by the RefSeq annotation (GRCh38 GFF file). A read is considered exactly aligned (green) if it is completely aligned, without mismatch, and starts and ends at the same chromosomal coordinates as the truth. A partial (light blue) must be within the true alignment. For misaligned reads (red), at least part of the alignment does not overlap the truth. Unmapped reads are shown in black. d.) The number of mismatches reflects mis-mappings, since by construction the iRefSeq exactly match the genome.

**Figure 7.**
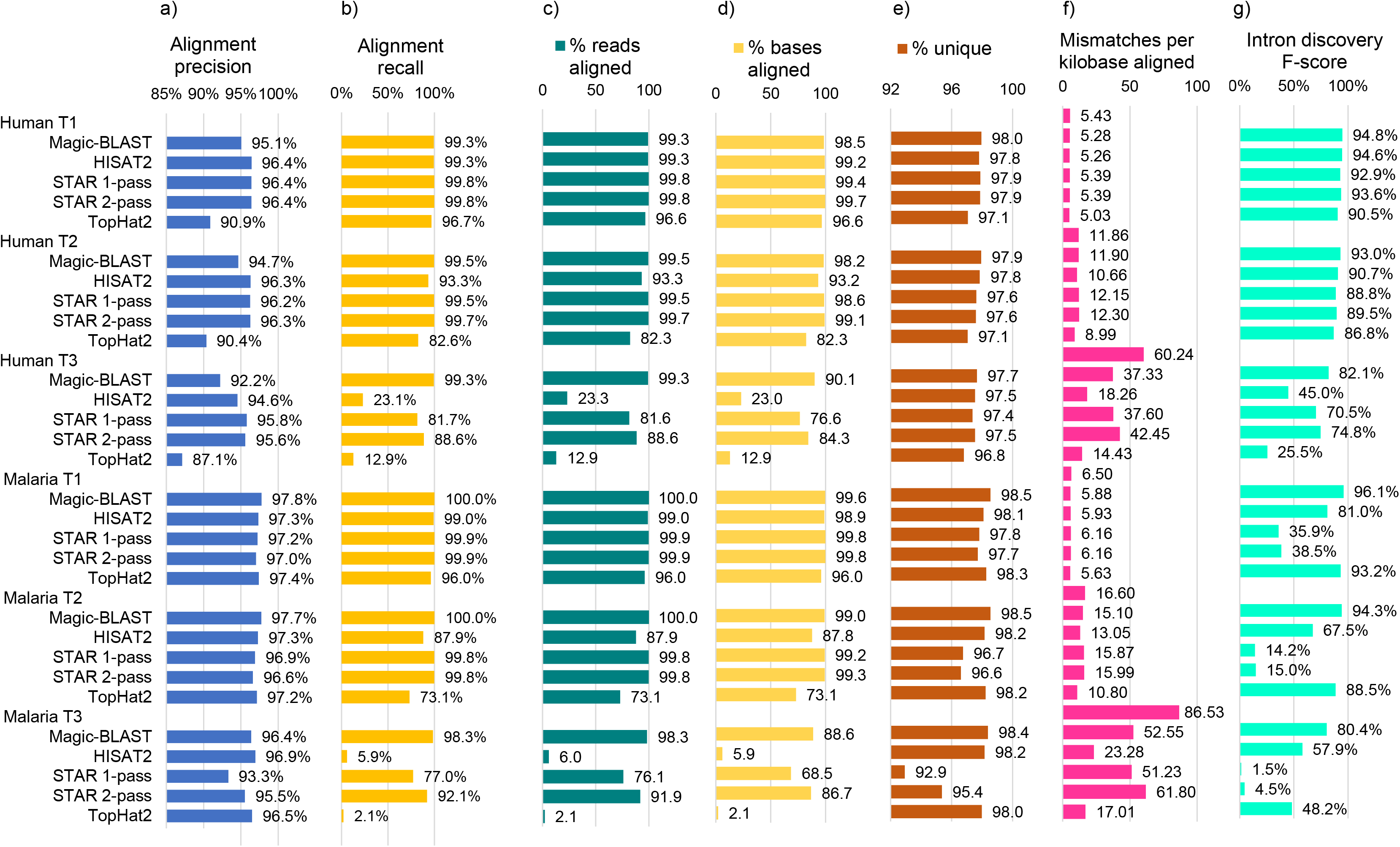
Performance of the aligners and quality control characteristics measured on the Baruzzo Illumina-like benchmark. First, the results of the comparison of all alignments to the truth are shown: a) alignment precision, b) alignment recall, counting as true an alignment which is identical or included in the truth (perfect or partial); each alignment for multiply aligned reads is counted. Second, the statistics of the alignment independently of the truth are shown: c) percentage of reads aligned. d) percentage of bases aligned. Note that the aligners differ on their view of what can be reported as an ‘alignment’ and on their thresholds for minimal aligned length and maximal level of mismatches. e) Percentage of reads aligned uniquely. A higher rate of unique alignments is desirable, but the true number of ambiguous reads is unknown. f) The number of mismatches per kilobase aligned. The true rate (first line of each dataset) was measured using the official benchmark mapping. A rate below the truth, correlated with lower alignment recall, characterizes an aligner which cannot deal with high levels of mismatches. g) the intron discovery F-score, measured for all introns (coverage at least 1). Magic-BLAST reaches the highest F-score for intron discovery in all datasets.

In all cases, even when no truth is available, we characterized the quality of the alignments by using the alignment statistics derived from the AliQC program (Supplementary S1.3), which includes the number of reads and bases aligned, unique versus multiple alignments, aligned length histograms, mismatches per read, type and position (Supplementary section 7 presents more information on the topics covered in this paragraph).

A simple and rich summary of the quality of the long reads alignments is provided in Figure 5 (and Figures S2), which shows for each aligner the histogram of aligned length as compared to the length of the reads. The best performance an aligner could possibly achieve would be to map each read along its entire length, so ideally the histogram of aligned length should be superimposed on the histogram of the read lengths (blue), especially for the longest reads (to the right). In the iRefSeq case of very long transcripts perfectly matching the genome (Figure 5a), TopHat2 fails, Magic-BLAST matches the read length histogram all the way from long to short transcripts. HISAT2 relaxed is better than HISAT2 and nearly as good as Magic-BLAST, but with and elbow of very short alignments. STAR long is distinctly lower over the longest reads. The situation is different for PacBio (Figure 5b) where the bulk of the sequences are between 1000 and 2700 bases long and show a high rate of sequencing errors, close to 20 mismatches per kb. There, STAR long finds some of the longest alignments but fails to align close to 30% of the reads. Magic-BLAST maps all reads, but often as partials because if an alignment presents a gap, Magic-BLAST reports only the highest scoring partial alignment. The curves for HISAT2 and HISAT2 relaxed are much lower, especially for long reads. In addition, HISAT2 relaxed creates a very large number of alignments shorter than 200 bases. The Roche reads are shorter, between 33 and 808 bases (Figure 5c), and some are now in the range acceptable to TopHat2, which aligns about half of the reads up to 600 bases. Alignment lengths for STAR long and Magic-BLAST are close to read lengths. The curve for HISAT2 is a little below. Both Magic-BLAST and HISAT2 relaxed produce a small number of shorter alignments, as is evident from the peaks on the very left end of the graph. The Illumina paired-end reads have a fixed length of 101+101 bases (Figure 5d). Ideally, all alignments should have 101 bases. The bar plot at the bottom shows that Magic-BLAST has the fewest complete alignments, but the situation is reversed for the longer Illumina reads 250+250 and 300+300 (Figure S2.2) where Magic-BLAST aligns 50% more bases than STAR and twice as many as HISAT2.

We now detail the results for the iRefSeq benchmark, then the six Baruzzo benchmarks, and finally the experimental runs from SRA.

The iRefSeq experiment evaluates the ability of the programs to align long spliced mRNA sequences exactly matching the genome, an idealized situation with no sequencing errors (Figure 6, Supplementary section 5). Magic-BLAST, HISAT2, HISAT2 relaxed, and STAR long produced results for this experiment. There is no bias, as Magic-BLAST was not used to prepare the RefSeq annotation.

Figure 6a displays the percentage of reads and bases aligned. Magic-BLAST performs the best. Similarly, at the intron level (Figure 6b), Magic-BLAST has more than 10 times fewer false positive introns than HISAT2 (158 versus 2336 or 2765) and four times fewer than STAR long. STAR long also misses the most introns (false negative) by a wide margin while Magic-BLAST misses the least.

Measuring the mapping accuracy by comparison to the iRefSeq annotation taken as the truth (Figure 6c) shows that 97.2% of the 45,108 iRefSeq mRNAs are perfectly mapped over their entire length by Magic-BLAST, 88.1% by HISAT2 relaxed, 84.1% by HISAT2, and 80.6% by STAR long (no clipped bases, substitutions or indels). There are also some correct but partial alignments. An important question is how often will a program misalign a read, since this would create noise in downstream analyses. This error happens 196 times (0.43%) with Magic-BLAST, but four to eight times more frequently in HISAT2 and STAR long. Most of the time, the misalignments are subtle, affecting just one or a few exons, but in 12 cases for Magic-BLAST, 103 for HISAT2 relaxed, 39 for HISAT2, and 100 for STAR long, the mRNA is wildly misaligned at a genomic site not overlapping the true position. Another functionally important case is the proper identification of the first and last exons, which reveal the location of promoters and 3’ ends regulatory regions. In Magic-BLAST, five alignments overlap the truth but extend outside of the annotated gene, creating a new or incorrect first or last exon, but this problem happens much more frequently in HISAT2 (191 and 79 times) and in STAR long (58 times). In a gene reconstruction project, this type of defect is hard to fix and may lead to incorrectly intertwined neighboring genes.

The iRefSeq mRNAs should match the genome exactly, and the number of mismatches shows how well each aligner performs (Figure 6d). The truth (blue) has no mismatch. Magic-BLAST has 771 mismatches, STAR long and HISAT2 relaxed 38018 and 66358 mismatches respectively, and STAR long aligns many fewer bases. On iRefSeq, Magic-BLAST alignments are superior to both HISAT2 and STAR long by all criteria.

We then assessed the accuracy of the alignments and measured the mapping statistics using 100+100 base Illumina-like simulated RNA-seq data from the Baruzzo benchmark (4) (Figure 7). The first two columns (7a and 7b) use the built-in truth to measure the precision and recall of the mapping: an alignment is counted as true if it overlaps the benchmark position, even if it is partial; otherwise it is counted as false (Supplementary section 6). In case of multiply aligned reads (Figure 7e), each alignment contributes. The human T1 set should be the easiest, and STAR, HISAT and Magic-BLAST do well, in this order: their precision is above 95% and recall above 99%. TopHat2 is noticeably lower. For the human T2 set, STAR 2-pass does the best, and HISAT2 and TopHat2 show some degradation. Magic-BLAST maintains a strong performance at the T3 level, STAR shows a significant degradation of the recall, and HISAT2 and TopHat2 perform very poorly. The percentage of reads and bases aligned tells a similar story (Figure 7c, 7d), with results degrading from T1 to T2 to T3, especially for HISAT2 and TopHat probably because the rate of mismatches in the benchmark, 60 per kb in Human T3 (Figure 7f), exceeds the thresholds of these programs.

The Plasmodium (malaria) benchmark sets should be more challenging owing to the AT-rich strong bias in the composition of the genome; also, the coverage is 100-fold deeper. The results are similar to the human benchmark; the main difference is that STAR is very slow and inefficient on malaria and produces huge numbers of false positive introns (Figure S3) and Magic-BLAST now has the best alignment precision and recall for T1, T2 and T3 (Figure 7).

Finally, we examine the ability of the programs to align experimental RNA-seq reads generated with different technologies. We find that, by all criteria, Magic-BLAST is the best mapper for all long reads, from Illumina 250+250 or 300+300 to Roche and PacBio (raw or edited) as well as for iRefSeq. The quality of the aligners is synthetized in Figure 8, which plots the number of matches against the number of mismatches summed over all ‘primary’ alignments (maximum one alignment per read). The mismatch rate depends on the sample and sequencing quality, its true level is unknown, but mis-mapping by the aligner will tend to increase the apparent rate. The most efficient aligner will align the largest number of bases – hence be most to the right - yet limit its mismatch rate to a minimum – hence be higher than other aligners with a similar number of matching bases. We indicate next to the best aligner and in the same color the number of mismatches per kilobase, which is our best estimate of the true level of mismatches in the sample (Figure 8 and S7.1). Magic-BLAST (red) has the largest number of matching bases in all cases where read length is larger than 101 bases. For Illumina 101+101, STAR 2-pass (green) aligns 1% more matching bases, but with a very high level of mismatches by Illumina standards, 9.3 per kb in STAR to be compared to 5.7 in Magic-BLAST. In general, the STAR family of programs tends to align with a much higher level of mismatches than Magic-BLAST, and as this is often coupled with a lower number of matches, one may suspect a high frequency of suboptimal mappings in STAR compared to Magic-BLAST. TopHat (light blue), and to a lesser extent HISAT2 (yellow) allow fewer mismatches hence stay near the left top corner.

**Figure 8:**
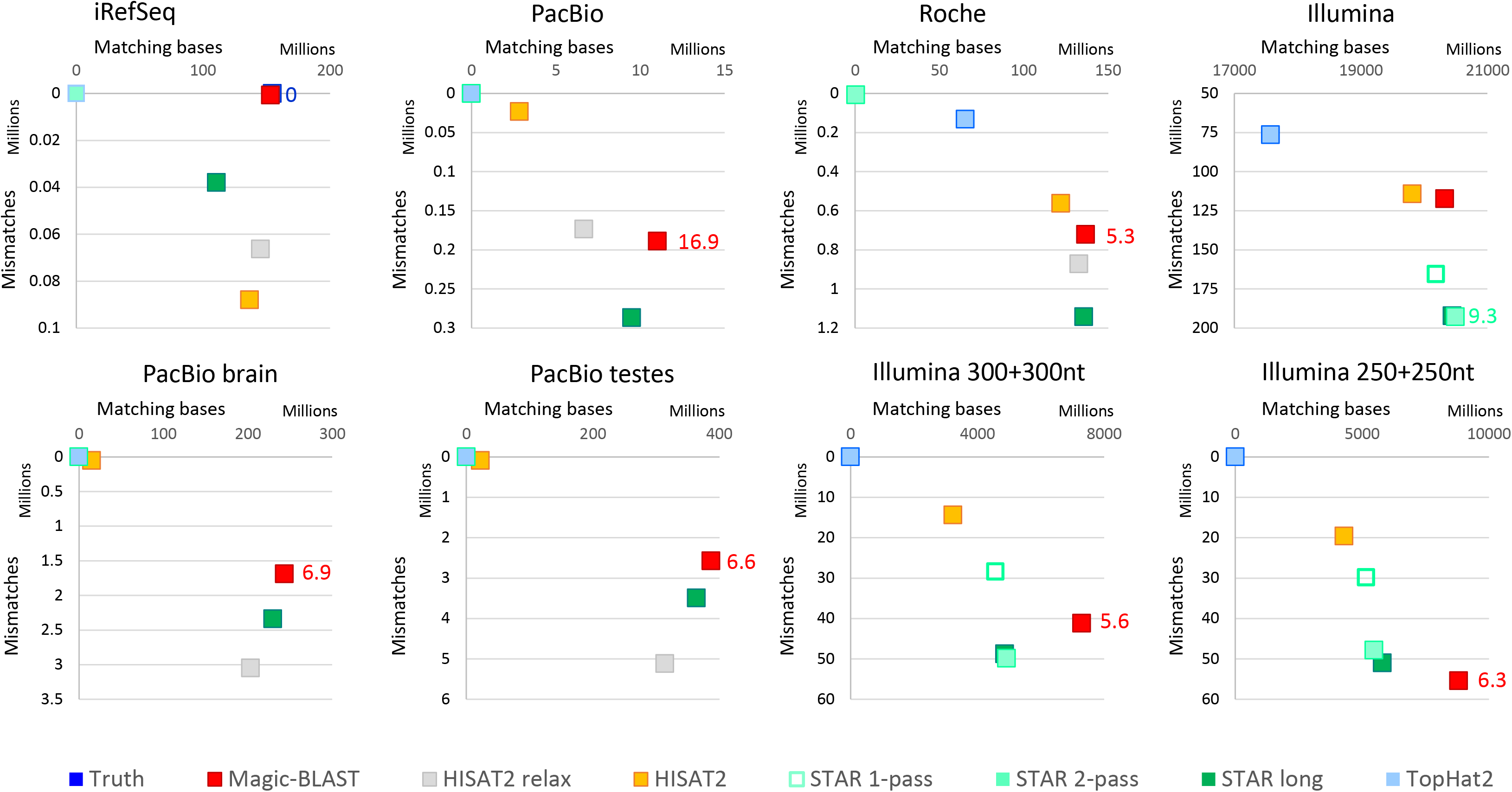
Number of matches (X-axis) versus mismatches (Y-Axis) for iRefSeq and seven experimental sets. The best aligner should be most to the right, then preferably more to the top, as mis-mappings will increase the mismatch rate and pull the aligner downwards. For each set, the number of mismatches per kilobase for the best aligner is shown in the color of the aligner, it likely reflects the actual mismatch rate in the sample. The iRefSeq sequences match the genome exactly, so the truth (dark blue square) is in the top right corner. The Magic-BLAST red square overlaps the truth. HISAT2 relaxed (grey) is the second most sensitive aligner (second most to the right) but has over 60,000 mismatches. STAR long has less mismatches but is by far the least sensitive. In all runs with long reads (3 PacBio, Roche and 2 long Illumina), Magic-BLAST has the most matching bases yet keeps a controlled mismatch rates. In Illumina 101+101, STAR long has slightly more matching bases but with a much higher mismatch rate. Notice that the PacBio brain and testes runs had been edited by the submitters using Illumina runs, which explains their low mismatch rate.

Figure 4 d-g presents some alignment statistics which brings more details and tell the same story. For longer Illumina reads (250+250 and 300+300 bases), Magic-BLAST aligns 50% more bases than STAR long or STAR and 70% more than HISAT2, and it does so while keeping a reasonable level of mismatches (5.6 to 6.3 per kb aligned). It also has the highest percentage of compatible pairs and remains the best at intron discovery (Supplementary sections 3, 4). TopHat2 results are lower by all criteria, except for intron discovery precision, good in very deep samples (malaria T1 T2 and Illumina) and in truncated 50+50 bases T1 pairs.

Supplementary sections 6 and 7 provide more details on all these aspects.

#### Run times

Figure 9 presents the CPU times and peak memory requirements for the alignments. A priori, the comparison seems straightforward, however, in presenting a fair assessment, the quality of the results matters. The table is colour coded by the percentage of matching bases relative to the best aligner for the dataset. For example, TopHat2 fails on PacBio (red) and only marginally succeeds on Roche (beige, 30 to 50% of top). Similarly, HISAT2 nearly fails on PacBio and Malaria T3 (brown, 5 to 10% of top). In such conditions, the speed of the code is less relevant. Looking at the colours in the table, Magic-BLAST is most often in the top aligner category (within 1% in 13 of 16 cases); it is less than 3% below the top in the three remaining cases, human benchmark T1 100+100 (-1.2%), T1 50+50 (-2.9%) and T2 (-1.1%). The second best aligner overall is STAR long 2-pass, which, as we discovered, can be used on any read length, but is slower on short reads than STAR 2-pass and less efficient than Magic-BLAST for long reads, from Illumina 250+250 paired ends to PacBio and iRefSeq. HISAT2 and Tophat2 rarely achieve top category (3 cases for HISAT2, 0 for HISAT2 relaxed and once for TopHat: T1 50+50).

**Figure 9.**
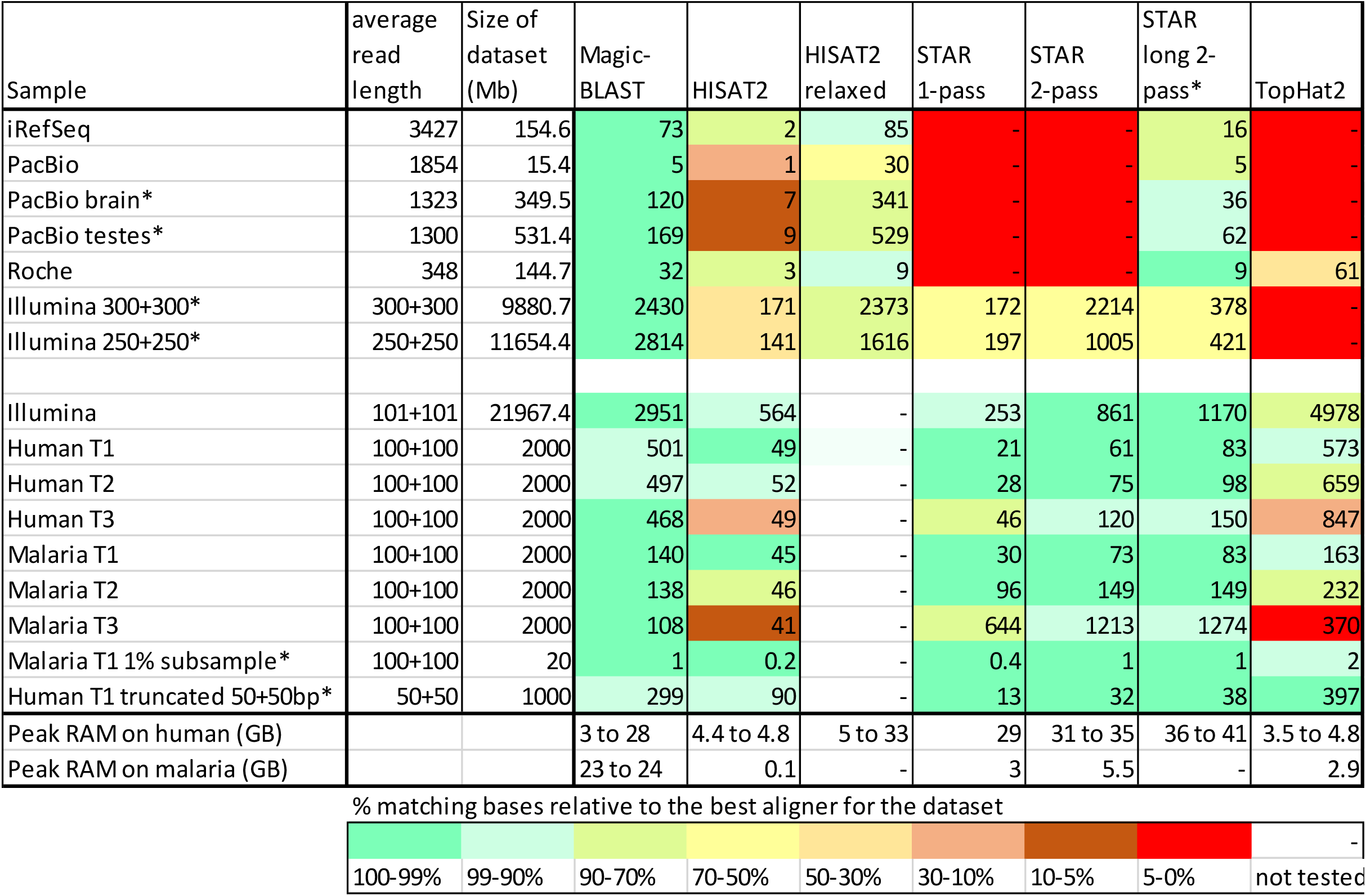
CPU (user plus system) time in minutes. The cells are color coded from dark green to red to show the percentage of matching bases relative to the best aligner(s) (dark green). Time is not given when there were no results. A blank cell indicates that the search was not performed. The last two rows show the range of peak memory usage for human and for malaria.

The memory requirement is also important (supplementary Table 1.2). HISAT2 uses by far the least memory (maximum 5 GB of memory in human, 0.1 GB in malaria) while STAR is the most greedy (27 to 42 GB). As a result, on an 8 core machines with 64 GB of RAM, one can run eight human samples in parallel using HISAT2, but only one using STAR long. Magic-BLAST uses slightly less memory than STAR on human but requires as much memory when aligning on malaria as on human. TopHat2, like HISAT2, generally uses little memory (less than 5 GB) except in human T1 50+50 (30 GB), the only case where it is a top aligner.

Consider now the speed of the programs and let us limit the comparison to good aligners (>90% of the top aligner, green and light green classes). On the benchmark human T1 or T2 reads, 50+50 and 100+100, STAR 2-pass or STAR long (and HISAT2 for T1) are the top aligners and the fastest codes, an order of magnitude faster than Magic-BLAST in human and three times faster in malaria. For the T3 cases which mimics cross-species mapping, Magic-BLAST is the only top aligner. Compared to STAR 2-pass, it is 3 times slower on human T3 and 10 times faster in malaria T3. STAR on malaria T3 consumed 24 times more CPU than on malaria T1 and 17 times more than on human T3. On the experimental Illumina 101+101, Magic-BLAST and STAR 2-pass are top aligners, and STAR is three times faster than Magic-BLAST. On datasets with longer reads, iRefSeq, PacBio, and Illumina 250 or 300, Magic-BLAST is the only top aligner, it is three to seven times slower than STAR long and one to three times faster than HISAT2 relaxed.

Most timings were performed on a 2.8 GHz Intel Xeon X5660 processor with 49 GB of RAM with a CentOS7 LINUX operating system. The time was measured with the LINUX time command by summing the reported user and system times. Before each run, the database and index files were cached in memory, to minimize influence of network and disk access on run time. A few runs, labelled with an asterisk, were timed later on a different machine which gave compatible results on runs tested on both machines. Supplementary section 1.1 discusses how to control the memory requirements when treating very large datasets with Magic-BLAST.

## DISCUSSION

We have examined the performance of four aligners with a wide variety of read lengths between 100 bases and 100 kilobases. First, we examined the performance of all programs with several experimental test sets from different platforms, with different lengths and characteristics. Second, we presented a new benchmark designed to test the ability of the programs to align very long sequences that have no mismatches to the genome. Finally, we looked at an artificial benchmark of short 100+100 base Illumina type reads, for human as well as a 100-fold deeper malaria runs, with three levels of mismatches. The aligners have different strengths and weaknesses, which reflect in part the strategic choices of the authors (e.g. favoring complete alignments or limiting the number of mismatches per read) and the characteristics of the implementations (e.g. second pass intron validation).

Magic-BLAST works for all datasets, produces better results on introns, long read alignments, and high-levels of mismatches, and is stable. It tries to align all reads, preferring a partial alignment to an unmapped read, because it knows how to precisely stops the alignment when encountering a lower matching area. As a consequence, some short alignments may be reported. For reads longer than 100 bases, it is a winning strategy. It aligns more bases and with less mismatches, and reliably discovers more introns. It is more exhaustive and more precise.

As demonstrated by the iRefSeq set, only Magic-BLAST, HISAT2 with non-default parameters and STAR long could align very long sequences, even if there were no mismatches. Both Magic-BLAST and HISAT2 relaxed were able to align all reads in the set while STAR long aligned 82%. In terms of bases aligned, Magic-BLAST found significantly more matching bases (99% versus 94% and 82%), and while we expect zero mismatches, Magic-BLAST had 68 to 130 times less mismatches than the other aligners. The same trends were found for the long experimental runs. On PacBio and long Illumina, Magic-BLAST aligned 70 to 76% of the bases while maintaining a low mismatch rate, the second-best aligned 65 to 68% of the bases.

For the shorter Illumina reads or the 100+100 simulated datasets with few mismatches (T1 or T2), other programs produced alignments comparable to or better than those of Magic-BLAST. For human T1 and T2, STAR 2-pass aligned the largest number of bases and reads and maintained an excellent alignment F-score. It degraded in T3 and in malaria, where Magic-BLAST maintained good performance. Magic-BLAST is not the best tool to align 50+50 paired end reads (Figure S7.1), where TopHat or the ten times faster but less efficient STAR would be better choices. HISAT2 was memory efficient and did rather well on short reads with few mismatches.

Intron discovery posed different challenges for the aligners. We looked at ROC curves, using the provided results for the benchmark sets and the annotations on the human genome as a guide for the experimental sets. As discussed, the representation of introns in the RefSeq annotation may be uneven. Certain tissue types may be underrepresented, while highly expressed genes from other tissues are certain to be included. This remark explains the need to measure the performance of an aligner on real data, with all the messiness of biology, but also on benchmark data with known results. It is clear from the ROC curves that our cautious approach to intron discovery pays dividends: Magic-BLAST is the best intron finder in all datasets, with far fewer false positives produced for a given number of true positives. The intron discovery F-score tells a similar story. Magic-BLAST has the best results in all datasets and its results are trustworthy, even at very low coverage. For the T1 or T2 human Illumina type reads with few mismatches, the difference in ROC curves between Magic-BLAST and HISAT2 was relatively small, but Magic-BLAST excelled for more distant matches or the compositionally biased malaria sets. For the T3 sets, Magic-BLAST had much better intron-finding F-scores than the other programs, consistent with its read mapping F-scores. For the iRefSeq, PacBio, Roche and long Illumina sequences, Magic-BLAST again produced the best introns results.

We also found that the intron discovery ROC curves for STAR 2-pass were worse than STAR 1-pass for deep sets such as the malaria sets or the Illumina 101+101, even though STAR 2-pass is expected to improve upon STAR 1-pass. For the Illumina run (zoom in Figure S3.1), the 1-pass curve lies just below Magic-BLAST, but the 2-pass curve is strongly shifted towards the unannotated/false positive. HISAT2, which also uses a 2-pass technique, is twice further than STAR 2-pass towards the noise. One could argue that most introns discovered by HISAT2 or STAR 2-pass in Illumina are real and missing from the annotation. However, our design contains seven controls where the true curve is vertical: in iRefSeq and in the Baruzzo sets, especially malaria, there is no doubt that the highly covered unannotated introns of STAR and HISAT2 are false positives, despite their high coverage. Both STAR 2-pass and HISAT2 perform better for the shallow T1 human set than the deep malaria T1. The optimal F-scores (Figure S4.1), calculated for the different aligners and experiments, are consistent with the ROC-curves in this regard. We have shown by subsampling that in both aligners, the second pass reinforces the false discoveries of the greedy first pass, and this tendency is also visible in all PacBio runs and Roche. Sadly, this type of noise cannot be erased, and if used in a gene reconstruction project, as was done in (17), those well supported but false positive introns will generate alternative splice variants that do not exist in the biological sample and will durably pollute reference gene models.

In Figure 9, we presented the run times for the various aligners and their relative performance on each dataset. For the longer sequences (first seven rows), from Illumina 250+250 to iRefSeq, Magic-BLAST was consistently in the best category. In some cases, Magic-BLAST could be 3-7 times slower than the next best aligner (STAR long or HISAT2 relaxed), but the quality of the results, both in terms of the alignments and intron finding, made it the best choice. For shorter sequences, such as Illumina 101+101 or the Baruzzo 100 base T1 and T2 benchmark sets, most aligners performed relatively well. Most were faster than Magic-BLAST and therefore a reasonable choice for short reads similar to the target. The other aligners did not perform as well for the more distant T3 Baruzzo sets and STAR slowed down dramatically for the T3 malaria case, becoming 10 times slower than Magic-BLAST.

Magic-BLAST was the only aligner to produce good results for a wide variety of sequence lengths, from 100 bases to 100 kilobases, compositions, or error rates without changes to the command-line whereas the standard options of other programs are often suboptimal (4). We also field-tested Magic-BLAST in several hackathons, allowing us to identify problems, hear user suggestions, and improve usability.

It is finally instructive to examine how the standard BLASTN algorithm handles spliced alignments. We search an mRNA (u00001.1) against the human genome reference assembly (GRCh38.p12) with BLASTN and find two problems. First, BLASTN identifies (apparently) strong matches on chromosomes 2, 14, 17, 20, 21, and 22 as well as two unplaced genomic scaffolds. A quick examination of the BLASTN alignments shows that all the matches are processed pseudogenes, except for the one on chromosome 17. Second, BLASTN does correctly identify the genomic exons on chromosome 17 but gives an imprecise result as shown in Figure 10. A spliced aligner, like Magic-BLAST, aligns this read only on chromosome 17 and correctly finds that the first exon ends at 85 on the query, and the second exon starts at 86 on the query. Additionally, BLASTN has no facility for recognizing paired reads so that it aligns and scores each read independently.

**Figure 10.**
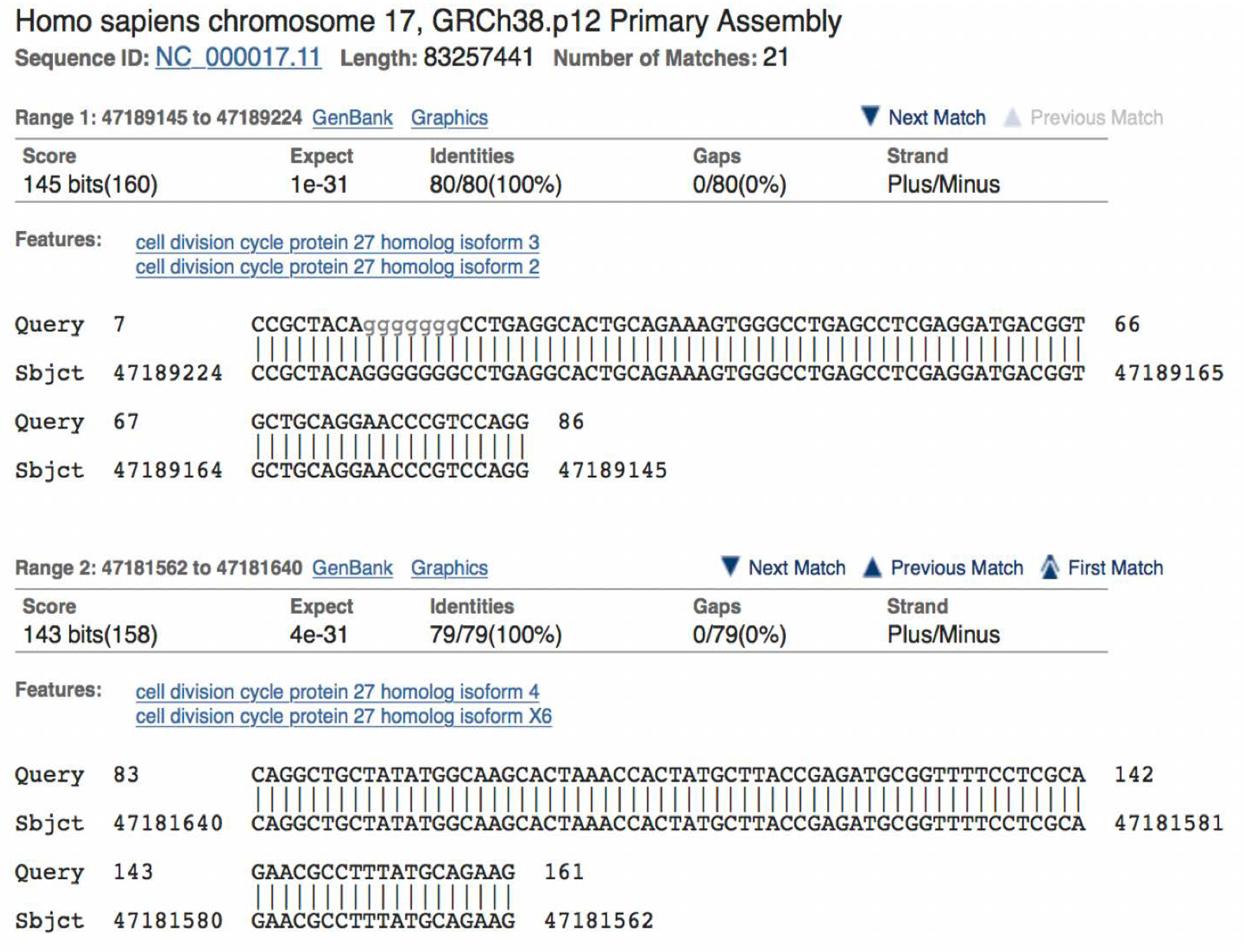
The first two BLASTN alignments of U00001 on human chromosome 17. U00001 is the query and is shown on the top of each row. The first exon really ends at base 85 of the mRNA, but BLASTN aligns the mRNA query to the first exon as well as an extra base in the intron. The second alignment starts three bases before the beginning of the second exon. BLASTN is not splice aware and aligns beyond the splice site, and is not aware of paired end sequencing.

### BLAST toolkit integration

Magic-BLAST takes full advantage of the BLAST Toolkit. It uses a BLAST database for reference sequences that compresses sequences 4-to-1, saving disk space and memory. The same database can also hold reference sequence metadata such as taxonomy, length, identifiers, and titles. The sequences and the metadata can be retrieved with the <u>blastdbcmd executable</u> (see https://www.ncbi.nlm.nih.gov/books/NBK279690/). Additionally, the databases can also hold user-supplied masking information for the reference sequences that can be selectively enabled. Similar to the BLAST+ programs, one can use the-seqid_list option to map sequences only to selected reference sequences in a larger BLAST database. There is a caveat though: not mapping the reads competitively on the entire genome will gather all the reads coming from the selected target, but also some from its cognate genes, homologous sequences and eventual pseudogenes.

For bioinformatics workflows, a significant advantage of Magic-BLAST is the flexibility in obtaining the sequence reads and the ease of setting up the reference set. It can map both RNA and DNA sequencing runs to a reference genome or transcriptome. Reference sequences can be given as FASTA, BLAST database, or as a list of NCBI accessions, but searches with a BLAST database are the fastest. Magic-BLAST has been used in several rapidly prototyped workflows (https://github.com/NCBI-Hackathons). Examples provided on project web pages show how users can download NCBI reference genomes, create BLAST databases and map SRA experiments with Magic-BLAST. For instance, Magic-BLAST was used for the fast estimation of the expression of a selected transcript in an SRA run (https://github.com/NCBI-Hackathons/deSRA). This project compared the expression of the transcripts in two NGS sets by quickly creating a small BLAST database for selected transcripts. Magic-BLAST then searched the two NGS projects against the BLAST database.

### Conclusion and next steps

We presented Magic-BLAST, a new tool for mapping next generation RNA-seq runs with read length between 100 nucleotides and 100 kilobases against a genome or a transcriptome. Its performance was compared with that of similar popular programs: HISAT2, STAR, and TopHat2. Magic-BLAST was the best intron finder on all tested sets, real or simulated, and the best aligner for all long reads, including Illumina 250+250 or 300+300, Roche 454, PacBio and full-length mRNA sequences.

Magic-BLAST integrates very well with other NCBI tools and services and is convenient to use since it recognizes NCBI accessions for SRA runs, mRNA or genomic sequences, and uses BLAST databases.

We are exploring ways to improve Magic-BLAST, such as exporting lists of introns and support, improving adapter detection, shortening the required exon length, and identifying repeats. We are also working closely with users to address their needs.

## AVAILABILITY

Magic-BLAST executables and source code are available at ftp://ftp.ncbi.nlm.nih.gov/blast/executables/magicblast/LATEST. Command line executables are provided for 64-bit Linux, Mac-OS and Windows systems. The package includes the makeblastdb program for creation of a BLAST database. The makeblastdb program is the same as distributed with the BLAST+ package and added here only for convenience. Basic operation instructions and examples are provided in the Magic-BLAST cookbook: https://ncbi.github.io/magicblast .

A master script used for the current analysis is available at https://github.com/ncbi/magicblast and described in the supplementary material section 1. All data come from external sources and can be downloaded by the master script, together with all our alignment SAM files (ftp://ftp.ncbi.nlm.nih.gov/blast/demo/magicblast_article).

## SUPPLEMENTARY DATA

Supplementary Data are available at NAR online. The supplement consists of a single document with seven sections, successively 1- Bioinformatics (aligners and analysis programs), 2- datasets and length histograms, 3- introns discovered per coverage (ROC curves), 4- introns precision and recall, 5- iRefSeq results, 6- alignment mapping precision and recall measured by comparison to the benchmark, 7- alignment statistics and quality control. Each section is associated with a large supplementary table, available in a single excel workbook. All comments are welcome.

## ACKNOWLEDGEMENT

We would like to thank David Lipman for proposing and encouraging this project and David Landsman for his insight and support. We would also like to thank Kim Pruitt, Richa Agarwala, Michael DiCuccio, Alex Morgulis, Terence Murphy, Eugene Yaschenko, Peter Cooper and Tao Tao for useful discussions, experiments, and feedback, the NCBI systems group for their expert help and Joe Meehan for his contribution to the AliQC python program.

